# T cell-derived IFNγ instructs ECM crosslinking by cardiac fibroblasts through LOXL3 in experimental cardiometabolic HFpEF

**DOI:** 10.64898/2026.03.16.712110

**Authors:** Ramona Emig, Zachary Robbe, Celina Kley, Sasha Smolgovsky, Joshua G Travers, Robert M Blanton, Timothy A McKinsey, Lauren D Black, Pilar Alcaide

## Abstract

**Background:** Heart failure with preserved ejection fraction (HFpEF) is a major clinical challenge characterized by diastolic dysfunction. Left ventricular stiffening and inflammation are hallmarks of HFpEF, yet the contribution of extracellular matrix (ECM) stiffness and the immune–stromal mechanisms driving ECM stiffening in cardiometabolic HFpEF remain poorly understood.

**Methods:** We used the murine “2-hit model” of cardiometabolic HFpEF, in which the combination of high fat diet and hypertension induced by L-NAME causes diastolic dysfunction. We evaluated diastolic function by echocardiography and ECM mechanics by uniaxial tensile testing of decellularized cardiac tissue. Functional *in vivo* studies included genetic depletion of T cells, interferon-γ (IFNγ) knockout mice, and pharmacological lysyl oxidase inhibition. We combined co-cultures of CD4^+^ T cells and cardiac fibroblasts (CFB) with mechanical testing of cardiac ECM and molecular biology to elucidate cellular and molecular mechanisms.

**Results:** Left ventricular ECM stiffness strongly correlated with impaired diastolic function in experimental cardiometabolic HFpEF. Cardiac CD4⁺ T cell infiltration was required for ECM stiffening and upregulation of lysyl oxidase enzymes in CFB. CD4^+^ T cell-derived IFNγ was both necessary and sufficient to induce LOXL3 in CFB, which increased ECM stiffness *in vitro*. Mechanistically, IFNγ signaling activated hypoxia-inducible factor-1α (HIF1α) in CFB, driving LOXL3 expression and subsequent collagen crosslinking. Genetic or pharmacologic disruption of this IFNγ–HIF1α–LOXL3 axis *in vivo* attenuated adverse ECM remodeling and improved diastolic function.

**Conclusions:** CD4⁺ T cells promote pathological ECM stiffening in cardiometabolic HFpEF through IFNγ-mediated, LOXL3-dependent ECM crosslinking by CFB. Targeting this immune–stromal pathway may offer a novel therapeutic strategy for HFpEF.

## Introduction

Heart failure with preserved ejection fraction (HFpEF) accounts for more than 50% of all heart failure cases and is strongly associated with cardiometabolic comorbidities such as obesity, hypertension, and diabetes (1). Despite its growing prevalence, the pathophysiological mechanisms underlying HFpEF remain incompletely understood, and effective therapies are lacking. A defining feature of HFpEF is diastolic dysfunction, which has been linked to myocardial extracellular matrix (ECM) remodeling and increased tissue stiffness (2). However, the cellular and molecular drivers of this maladaptive mechanical remodeling in the context of cardiometabolic disease remain elusive.

Emerging evidence suggests that chronic low-grade inflammation contributes to HFpEF pathogenesis (3,4), yet the specific immune cell subsets and their interactions with cardiac stromal cells have not been fully delineated. CD4⁺ T cells, which are enriched in the myocardium during cardiometabolic stress induced by obesity and hypertension (5), have been implicated in fibrotic remodeling through crosstalk with cardiac fibroblasts (CFB) in other etiologies of HF (6,7). CFB are the primary cell type maintaining and remodeling the ECM. They exhibit profound, dynamic phenotypic changes in response to mechanical and biochemical cues, including upregulation of enzymes that mediate collagen crosslinking and tissue stiffening. Among these, lysyl oxidase family members have emerged as critical regulators of ECM mechanics through collagen crosslinking, yet their role in cardiometabolic HFpEF has not been defined (8–10). Whether T cell interactions with CFB increase the expression of ECM crosslinking enzymes and thereby promote myocardial stiffening is unknown.

Here, using a murine model of cardiometabolic HFpEF, we measured cardiac stiffening and ECM mechanical remodeling using uniaxial tensile testing and performed a combination of immunological, pharmacological and physiological approaches. We demonstrate that elevated left ventricular (LV) ECM stiffness in cardiometabolic HFpEF is driven by CD4^+^ T cells through induction of lysyl oxidase family member lysyl oxidase-like 3 (LOXL3) in CFB. We find that interferon-γ (IFNγ) activates hypoxia-inducible factor-1α (HIF1α) signaling and is critical for LOXL3 mediated ECM crosslinking by CFB. Genetic and pharmacologic perturbation of this pathway *in vivo* attenuates ECM stiffening and improves diastolic function.

## Methods

### Animal models

Male wild-type (WT) (C57Bl/6J, 000664), *Tcra^−/−^* (B6.129S2-Tcra^tm1Mom^/J, 002116) or *Ifng^−/−^*(B6.129S7-*Ifng^tm1Ts^*/J, 002287, all from Jackson Labs) mice were 8-12 weeks old at start of experiments. Mice were given a high fat diet (HFD) with 60% of caloric intake from lard (Research Diets Inc. D12492) and 0.5 g/L of the hypertension inducing agent L-nitro-arginine-methyl-esther (L-NAME, SigmaAldrich N5751) in drinking water for 3 weeks or 5 weeks as indicated. Control groups received standard chow and drinking water (STD). A subpopulation of WT mice received daily intra-peritoneal injections of the pan-lysyl oxidase inhibitor β-aminopropionitrile (BAPN, SigmaAldrich A3134, dissolved in PBS) of 100 mg/kg body weight from week 3-5 of HFD+L-NAME treatment. Other mice were subjected to intra-peritoneal injections of recombinant IFNγ (25 kU/injection in 100 µL PBS, Peprotech, 315-05). Control mice for BAPN and IFNγ treated mice received PBS injections of equivalent volume. Mice had *ad libitum* access to food and water and were kept at 12h day-night cycle. All animal experiments were approved by the local authorities and carried out in compliance with Institutional Animal Care and Use Committee (IACUC) requirements.

### Echocardiography

Systolic and diastolic cardiac function were assessed using transthoracic echocardiography in anesthetized mice as previously described (7). Mice were kept on a heated stage in supine position, with heart and respiratory rates continuously monitored *via* stage electrodes. Heart rate was kept between 450 and 550 bpm. Depilatory cream (Nair) was used to remove fur on the chest, and ultrasonic coupling gel was applied onto the chest for imaging with a 22-55 MHz echocardiography transducer (MS550D; Vevo 2100, FUJIFILM VisualSonics). Once the LV was clearly visualized in short axis view, LV end-systolic and end-diastolic dimensions (M-mode) were measured, and the LV ejection fraction was calculated. LV geometry was assessed using LV weights, LV anterior and posterior wall thickness, and end-diastolic volume. Parameters for diastolic function, including the ratio of early-to-late mitral valve inflow velocity (E/A) were derived from pulsed wave Doppler of transmitral flow in an apical four chamber view. Parameters of cardiac systolic and diastolic function were measured by averaging of values obtained from 8 cardiac cycles.

### Cardiac tissue decellularization and mechanical testing

Transverse sections of the LV of 1-2 mm thickness were incubated in decellularization buffer (1% w/v sodium-dodecyl sulfate, 1% v/v Triton-X100 in PBS) for 3-5 days at room temperature on a rotator until the tissue was translucent. After washing in PBS, the decellularized tissue was submerged in PBS with 0.05% w/v sodium azide for storage.

For mechanical testing, a 1-3 mm long strip corresponding to the LV free wall was dissected from the decellularized tissue and mounted on a custom uniaxial tensile testing system (11). On one end, it was glued to a static lever, while the other end was fixed to a Dual Mode Lever System with a 1 N load cell (AuroraScientic #6350*358). For control and data digitization, the system was coupled to a digital input/output instrument (National Instruments USB-6221). Using an in-house LabVIEW script (LabVIEW 2011, National Instruments) uniaxial tension was applied in displacement control mode. Output was recorded at 30 samples/s. Before starting a strain protocol, the membrane was brought into a straight planar position with 5 mN pre-load applied. Then, a cyclic strain protocol was executed and at least 20 cycles were recorded. During measurements, the tissue was submerged in saline. Analysis of the stress-strain data was performed using a custom MatLab script (MatLab 2025b). Briefly, the recorded force and displacement were converted into stress and strain, respectively. Then, a linear relation was computed for the stress-strain relation during tissue stretch (not relaxation) at 10-15% strain. The slope of the linear relation is referred to as Elastic modulus.

To assess the effects of the CFB secretome on ECM mechanical properties, right ventricular tissue was harvested from 6–12-week-old C57Bl/6J mice and decellularized as above. Decellularized tissues preparations (dECM) were subjected to uniaxial tensile testing, then sterilized using ethanol, and incubated in conditioned media from CFB for 16-20h before repeating the tensile test. CFB conditioned media was supplemented with the lysyl oxidase inhibitor BAPN (500 µM, SigmaAldrich A3134) where indicated. Data was analyzed as above and displayed as fold-change of the Elastic modulus between before and after the incubation.

### Picrosirius Red staining

For assessment of LV fibrosis, one third of the LV was fixed in 10% formalin, embedded in paraffin, sectioned (5 µm thickness) and mounted onto microscopy slides. After deparaffinization, the slides were stained in Picrosirius Red staining solution (1 g/L Direct Red 80, SigmaAldrich 365548) for 60 min followed by two washing steps in acidified water. The slides were then dehydrated and mounted using non-aqueous mounting medium (Depex, EMS 13514). Per heart, five representative fields of view not containing vessels were imaged, and the collagen area fraction was quantified using FIJI (12). Each data point represents the average of five images from the same section.

### ECM quantification

To quantitatively assess ECM content in LV tissue, insoluble matrix proteins were enriched from ∼20 mg of frozen cardiac tissue using the Compartment Protein Extraction Kit (EMD Millipore 2145) according to the manufacturer’s protocol. The remaining insoluble pellet was washed twice in PBS containing protease inhibitors. Pellets were dried overnight at room temperature and dry pellet weight was normalized to the weight of the input tissue.

### Heart digestion for flow cytometric analysis

LV myocardium was harvested from anesthetized mice after terminal blood draw, finely minced using a razor blade and digested using 0.895 mg/mL of Collagenase Type-II (Gibco 17101015) in phosphate buffered saline (PBS) to achieve a single cell suspension. After 20 min, the tissue was dissociated mechanically using a 19 Gauge stainless steel cannula. After a total digestion time of 30 min, the suspension was filtered through a 100 µm cell strainer to remove remaining tissue chunks. The resulting single cell suspension was then stained for flow cytometry using fluorophore coupled antibodies (Table S1) at the indicated dilutions in FACS buffer (PBS +2% heat-inactivated fetal bovine serum) at 4 C in the dark. After 20 min, the cells were washed in FACS buffer, and 50 μL/sample Precision Count Beads (BioLegend 424902) were added to quantify absolute cell numbers. Spectral flow cytometry was performed on the Cytek® Aurora flow cytometer. After spectral unmixing, flow cytometry data was analyzed using FloJo (BD BioScience v10.10.0).

### Human bulk RNA sequencing analysis

Hahn *et al* performed bulk RNA sequencing on LV septum biopsies from control patients and patients with HFpEF (13). We downloaded the list of differentially expressed genes and extracted those with significant differences (adjusted p<0.05) between healthy controls and HFpEF patients. Genes with significantly higher expression in HFpEF patients were subjected to GO term analysis. Then, raw reads per patient were downloaded and data from control patients as well as patients in HFpEF was extracted. Using an in-house Python script generated with support from Microsoft Co-pilot, we computed Pearson’s correlation coefficient (r) and p-value for the correlation between *CD4* and all collagen or lysyl oxidase encoding genes.

### Single cell RNA sequencing analysis

We downloaded raw single cell RNA sequencing data from a publicly available source (14). Count data was processed sample wise with application of the following filter cut-offs: >200 Features, <25% mitochondrial genes, <1% ribosomal genes and >500 RNA counts using Seurat version 5.3.1. Data was log-normalized, and samples were clustered by principal component analysis using 30 dimensions followed by FindCluster function within the Seurat package with a resolution of 0.7. Cell type markers were calculated using the FindMarkers function with default parameters (Wilcoxon test) in Seurat and cell types were manually annotated based on known canonical markers. CFB were subsetted and re-clustered by principal component analysis using 30 dimensions and the FindCluster function with a resolution of 0.5. Percent contribution to the CFB population for each cluster was calculated based on the number of single cells in each cluster in control and HFpEF groups, respectively. Subcluster markers were calculated using the FindMarkers function with default parameters. Significant marker genes of cluster 0 (adjusted p-value<0.05) were subjected to GO analysis using Panther. Lastly, a lysyl oxidase score was calculated as the sum of the expression of all lysyl oxidase family members (*Lox*, *Loxl1*, *Loxl2*, *Loxl3*, *Loxl4*) per cell.

### LOXL3 ELISA

For protein analyses, snap-frozen LV tissue samples were thawed on ice and mechanically disrupted in 100 μL RIPA buffer (ThermoFisher J62524.AE) containing protease (ThermoFisher A32953) and phosphatase inhibitors (ThermoFisher A32957). The lysates were cleared from debris by centrifugation (10,000 rpm, 5 min, 4 C) and total protein content was determined using a bicinchoninic acid assay (ThermoFisher 23225) according to the manufacturer’s instructions. Lysates were diluted to 10 μg/mL and applied to enzyme-linked immunosorbent assay targeting LOXL3 (LS Bioscience LS-F14541-1).

### Splenic CD4^+^ T cell isolation, culture and polarization

Splenic CD4^+^ T cells were isolated by positive selection using CD4^+^ magnetic beads (Miltenyi Biotec 130-117-043) from spleens of 7–12-week-old C57Bl/6J mice according to previously established protocols (15,16). As basal T cell culture media we used RPMI supplemented with 10% (v/v) heat-inactivated fetal bovine serum (Gemini Bio S11550), 1.2 mM sodium pyruvate (Gibco, 11360-070), 0.1% (w/v) sodium bicarbonate (Gibco 25080-094), 1x GlutaMAX (Gibco A12860-01), 1x Pencillin/Streptomycin (Gibco 15070-063) and 0.0005 (v/v) β-mercaptoethanol (Sigma M3148). For *in vitro* experiments, cells were cultured at 2 million cells per mL in the presence of plate-bound αCD3 (coating with 2.5 μg/mL, BioLegend 100253) and soluble αCD28 (1 μg/mL, BioLegend 102102) and IL-2 (25 U/mL, Peprotech 212-12) for 3 days at 37°C to generate activated CD4^+^ T cell blasts. After 3 days, cells were expanded at 1:2 in media containing IL-2 (25 U/mL). After 24h, cells were centrifuged at 10,000 g for 5 min at room temperature and supernatants were collected under sterile conditions. These conditioned media were used to treat CFB as described below.

### CFB isolation, culture and treatments

To isolate CFB, LV of 4–8-week-old WT C57Bl/6J mice were minced and digested using 2 mg/mL Collagenase Type I (Gibco 17100017) for 25 min (including mechanical disruption by cannulation at 20 min digestion time). The suspension was filtered through a 100 µm cell strainer and plated onto gelatine-coated tissue culture plates. At 90% confluency, CFB were detached using Trypsin-EDTA (Gibco 25300054) and expanded. Experiments were performed at passage 2 after isolation. For RNA isolation, CFB were plated at 50,000 cells/well in 12 well plates. Basic CFB culture media was DMEM supplemented with 10% (v/v) fetal bovine serum (Gemini Bio S11550), 1% (v/) Insulin-Transferrin-Selenium (Gibco 41400-045) and 1% (v/v) Penicillin/Streptomycin (Gibco 15070-063). For immunofluorescence, CFB were plated at 5,000 cells/well onto gelatin-coated glass-coverslips in 24 well plates. 24h after plating, cells were starved in culture media containing 2% fetal bovine serum (FBS) for another 24h. Then, the indicated treatments were applied as follows: conditioned media from activated CD4^+^ T cell blasts (1:4 in CFB culture media), recombinant interferon-γ (IFNγ, 100 U/mL, Peprotech, 315-05), recombinant IL-2 (25 U/mL), recombinant IL-4 (500 ng/mL, Peprotech 214-14), recombinant transforming growth factor-β (TGFβ, 100 ng/mL, Peprotech 100-21). In selected experiments, CFB were additionally treated with the following inhibitors: αIFNγ (10 ug/mL, BioLegend 505834), Echinomycin (HIF-1α inhibitor, 5 nM, Tocris Bioscience 5520).

### qPCR

RNA from cultured CFB was isolated using the RNeasy kit (Qiagen 74106) according to the manufacturer’s instructions and RNA yield was assessed spectrometrically. Adjusted amounts of RNA were reverse transcribed into complementary DNA (cDNA) using High-Capacity cDNA kit (Fisher Scientific 43-874-06). A total of 9 ng/reaction was then subjected to quantitative real-time polymerase chain reaction using the primer sequences in Table S2. Expression of target genes was normalized by the expression of the reference gene *Rpl19* and is expressed as fold change with respect to the untreated control condition.

### Collagen contraction assay

To assess their contractile properties, primary murine CFB were cultured in a neutral Type1 Collagen solution (1 mg/mL, Advanced Biomatrix 5074) at 150,000 cells in 500 µL per well of a 24 well plate. After an initial solidification period of 1h, 600 µL of CFB culture media with the indicated treatments were added to each well followed by immediate release of the collagen hydrogel from the edges of the well. Images of the collagen disks were taken after 24 h. Disk area was quantified from these images and expressed as percentage of the area of the entire well.

### Immunofluorescence

CFB cultured on glass coverslips and treated as indicated were chemically fixed using 4% paraformaldehyde for 10 min. After permeabilization with 0.1% Triton-X100 in PBS (15 min) and blocking in 5% normal goat serum in PBS (1h), primary antibodies were applied overnight at 4 C. Appropriate secondary antibodies were applied for 1h at room temperature. Coverslips were mounted onto microscopy slides using Permafluor mounting medium containing DAPI (Southern Biotech 0100-20) for nuclear counterstain. Cells were imaged within 48 h of staining. For quantification of smooth muscle actin (SMA) and collagen-I (COL1A1), the mean fluorescence intensity across the entire image was averaged. For quantification of nuclear HIF1α, a region of interest was defined based on the nuclear signal and the average intensity of the HIF1α signal in this region was computed. Five representative images were acquired per coverslip, and each data point represents the average of those.

### Statistical analysis

Statistical analysis was performed using GraphPad Prism 10. All data are presented as mean±standard error of the mean. Individual data points represent data from individual mice (for *in vivo* studies) or independent biological replicates (for *in vitro* studies). Normal distribution was assessed using Shapiro-Wilk test. Two group comparisons were done by unpaired Student’s t-test for normally distributed data or Mann-Whitney test for non-normally distributed data. Multiple group comparisons were done using 1way or 2way ANOVA with Tukey’s multiple comparison test as appropriate. Differences were considered statistically significant at p<0.05.

## Results

### Mechanical ECM remodeling correlates with diastolic dysfunction in experimental cardiometabolic HFpEF

We subjected 8-12-week-old WT mice to the combination of HFD and L-NAME for 5 weeks which induced diastolic dysfunction, while ejection fraction was preserved (Figure 1A, Table S3). Doppler echocardiography confirmed a higher ratio of early/late mitral valve inflow velocity which is indicative of diastolic dysfunction, as expected based on recent publications (Figure 1B and 1C) (5,17). As diastolic function depends on the extensibility of the ECM of the LV, we next aimed to assess the mechanical properties of the LV ECM. To this end, we prepared transverse sections of the LV free wall and subjected them to decellularization. We then assessed the elastic modulus as a measure of stiffness of the ECM (Figure 1D and 1E). The LV-ECM derived from the hearts of mice treated with HFD+L-NAME was significantly higher than that of mice receiving STD chow (Figure 1F), while we did not detect significant differences in the stiffness of the right ventricular ECM between both groups (Fig S1A). Importantly, only the combination of HFD and L-NAME treatment, but not either treatment alone, caused high LV-ECM stiffness after 5 weeks (Figure 1G). Importantly, the stiffness of the LV-ECM correlated significantly with the E/A ratio across STD and H/L groups (Pearson’s R^2^: 0.512, p value: 0.009), highlighting the importance of mechanical ECM remodeling for diastolic function (Figure 1H). Noteworthy, the higher stiffness of the LV-ECM was not associated with significant collagen deposition that could be detected by picrosirius red staining (Figure 1I). Similarly, we did not find significant differences in the abundance of insoluble ECM proteins (expressed percent of insoluble ECM weight over tissue weight, Figure 1J). Flow cytometry results showed no significant differences in total CD45^−^CD31^−^MESFK4^+^ CFB numbers (Figure S1B and S1C). Together, these data demonstrate significant stiffening of the LV-ECM in response to HFD/L-NAME treatment without increasing ECM deposition, and suggests that mechanical remodeling of the ECM, rather than ECM abundance, is a critical determinant of diastolic dysfunction in response to HFD/LNAME.

**Figure 1:**
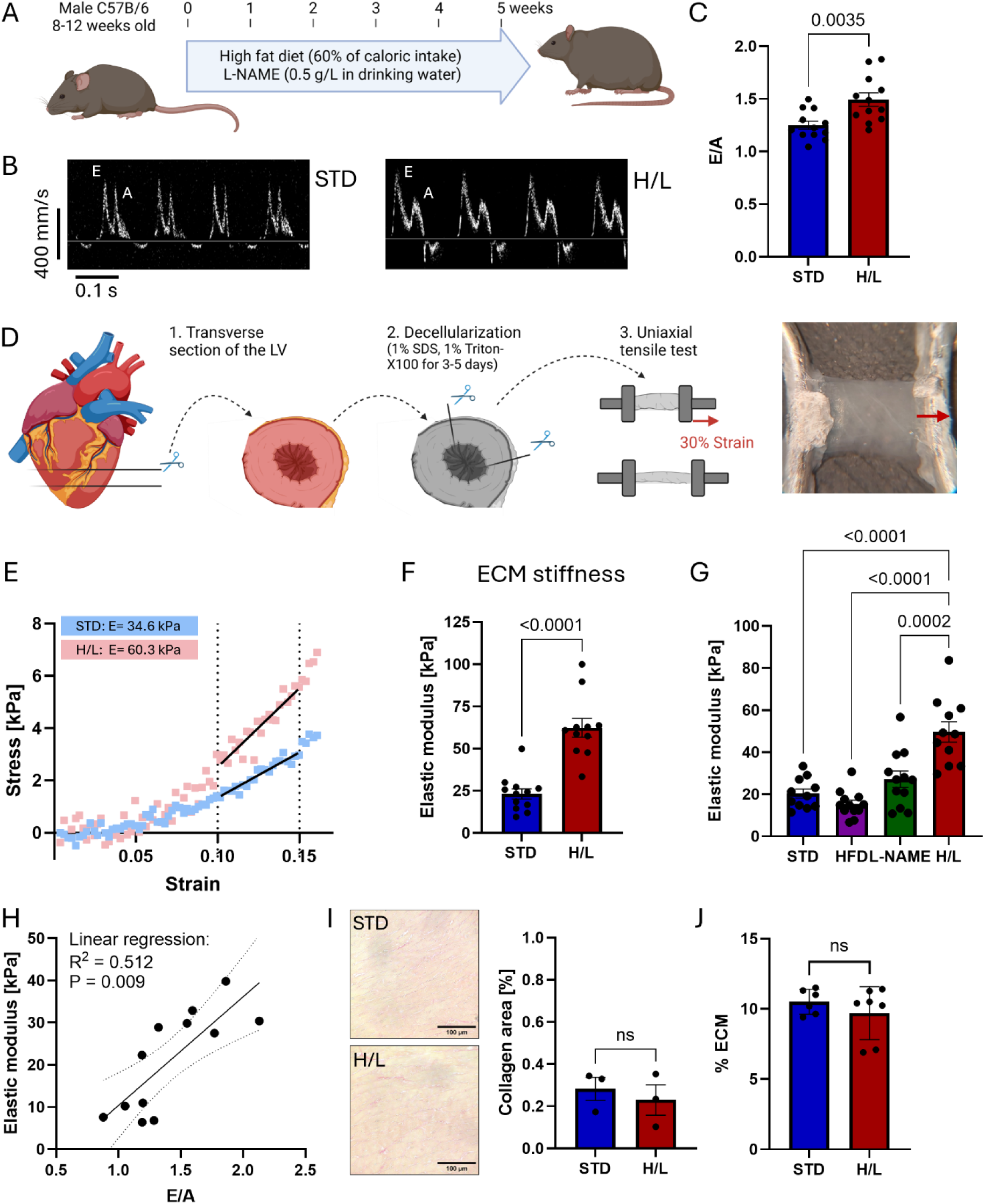
Mechanical ECM remodeling correlates with diastolic dysfunction in cardiometabolic HFpEF. A: 8-12-week-old wild-type C57Bl/6J mice received standard chow and water (STD) or the combination of high fat diet (HFD, 60% fat) and drinking water containing 0.5 g/L of the hypertension-inducing agent L-nitro-arginine-methyl-ester (L-NAME) for 5 weeks. B+C: Diastolic function assessed by Doppler-mode echocardiography: representative recordings of mitral valve inflow velocity (B) and ratio of early/late mitral valve inflow velocity (E/A, N=11-12/group, C). D: Preparation of decellularized left ventricular (LV) ECM samples for uniaxial tensile test. E: Representative recordings of uniaxial tensile test on LV-ECM samples from STD vs H/L-treated mice. F: Elastic modulus of LV-ECM (N=12/group). G: Elastic modulus of LV-ECM from mice on STD diet, HFD, L-NAME in drinking water, or the combination of HFD and L-NAME (H/L) for 5 weeks. H: Simple linear regression between E/A and LV-ECM Elastic modulus after 5 weeks of STD- or H/L-treatment. I: Representative images of Picrosirius red staining of LV sections of STD- or H/L-treated mice with quantification of collagen area fraction (N=3/group). J: Weight of dry, insoluble ECM peptides as a fraction of whole tissue weight for LV from STD- vs H/L-treated mice (N=6-7/group).

### Mechanical ECM remodeling in cardiometabolic HFpEF is T cell-dependent

We previously demonstrated that HFD/L-NAME causes a cardiotropic response in inflammatory CD4+ T cells and that T cell-deficient mice are protected from diastolic dysfunction in this model (5). Thus, we investigated the temporal relationship between cardiac CD4^+^ T cell infiltration and ECM stiffening. Using flow cytometry to identify CD4^+^ T cells in the LV of mice treated with STD chow or HFD+L-NAME for 3 or 5 weeks (Figure S2), we found increased LV CD4^+^ T cell numbers at 5 weeks compared to STD fed controls, while no significant differences were observed at 3 weeks. (Fig 2A). Remarkably, the time course of increase in LV-ECM stiffness correlated with CD4^+^ T cell infiltration. Specifically, LV-ECM stiffness was significantly higher after 5 weeks of HFD/L-NAME compared to STD mice, while no significant differences were identified at 3 weeks of HFD/L-NAME, a time point at which CD4^+^ T cells are not yet significantly increased in the heart (Figure 2B). To assess if the increase in CD4^+^ T cells in the hearts was causally related to the stiffness of the ECM and diastolic function, we treated T cell deficient mice (*Tcra^−/−^*) alongside WT mice with HFD/L-NAME or STD chow for 5 weeks. *Tcra^−/−^* mice did not show significant diastolic dysfunction in response to HFD/L-NAME treatment (Figure 2C and 2D, Table S4). Strikingly, LV-ECM stiffness from HFD/L-NAME treated *Tcra^−/−^* mice did not increase compared to STD. In fact, LV-ECM stiffness was significantly lower in HFD/L-LAME treated *Tcra^−/−^* mice compared to WT mice (Figure 2E and 2F). These data confirm the importance of CD4^+^ T cells for diastolic dysfunction and support that CD4^+^ T cells are functionally involved in ECM stiffening in response to HFD/L-NAME.

**Figure 2:**
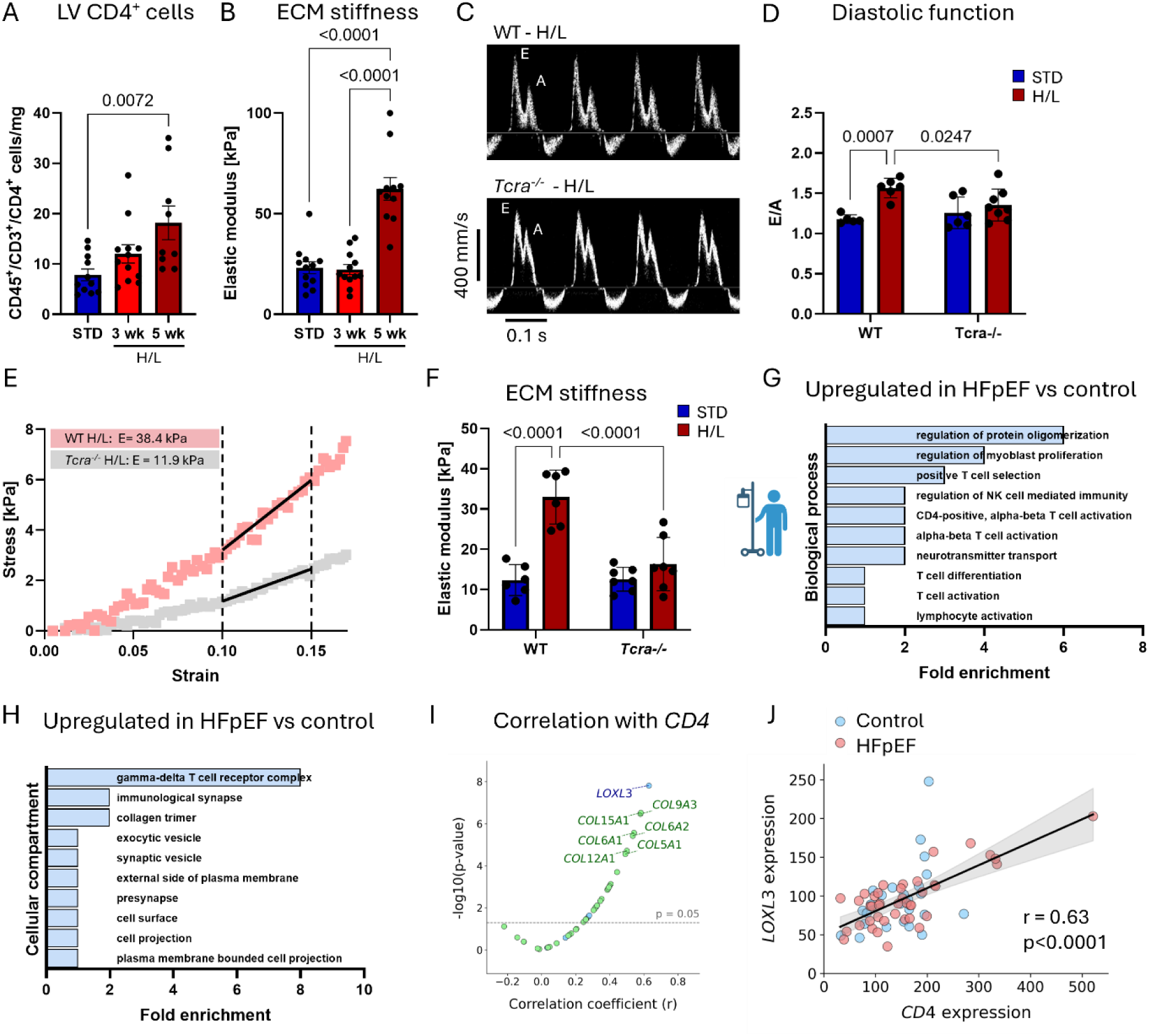
Mechanical ECM remodeling in murine cardiometabolic HFpEF is T cell-dependent and human HFpEF transcriptome is characterized by a T cell signature. A: Number of CD4^+^ T cells identified by flow cytometry in mice fed a STD diet for 3 weeks or H/L for 3 or 5 weeks (N=12/group). B: Elastic modulus of LV-ECM of STD- (5 weeks) or H/L-treated mice (3 or 5 weeks, N=12/group). C+D: Diastolic function assessed by Doppler-mode echocardiography in wild-type or *Tcra^−/−^*C57Bl/6J mice after 5 weeks of STD or H/L treatment: representative recordings of mitral valve inflow velocity (C) and ratio of early/late mitral valve inflow velocity (E/A, D, N=6-8/group). E+F: Uniaxial tensile testing of decellularized LV-ECM derived from WT or *Tcra^−/−^* mice after 5 weeks of STD- or H/L-treatment: Representative recordings (E) and Elastic modulus (F, N=6-8/group). G+H: Bulk RNA sequencing of human LV tissue from control and HFpEF patients. GO analysis of significantly upregulated genes in HFpEF compared to control samples. Shown are the 10 most significantly enriched biological processes (G) and cellular compartments (H). I: Correlation analysis of collagen (green) and lysyl oxidase (blue) encoding genes with *CD4*. J: Simple linear regression and 95% confidence interval between gene expression of the lysyl oxidase *LOXL3* and *CD4* across control and HFpEF samples.

Given the important role of T cells in murine cardiometabolic HFpEF, we next asked if similar features were present in human HFpEF patients using a publicly available bulk RNA sequencing data set from human HFpEF patients and healthy control patients (13). We found that terms related to (CD4^+^) T cell activation and differentiation were significantly overrepresented within genes upregulated in HFpEF patients (Figure 2G). We also found the cellular compartment *Collagen trimer* to be overrepresented in HFpEF patients suggesting that collagen crosslinking might be involved in HFpEF (Figure 2H). To examine a potential relation between the presence of CD4^+^ T cells and collagen remodeling, we compared the correlation coefficient and p-value of all collagen and lysyl oxidase genes, central to collagen crosslinking and ECM remodeling, with *CD4* gene expression, a proxy for the number of CD4^+^ T cells in the tissue. This analysis identified the strongest and most significant correlation occurs between *CD4* and the lysyl oxidase family member *LOXL3* (Pearson’s R: 0.63, p value: <0.0001, Figure 2I). The correlation between *CD4* and *LOXL3* was statistically significant across patient groups (Figure 2J). Thus, we hypothesized that lysyl oxidation by LOXL3 might represent a mechanism by which the stiffness of existing ECM is increased in the absence of additional ECM production.

### CFB express higher levels of lysyl oxidases in cardiometabolic HFpEF

We next investigated the cellular source of lysyl oxidases in the myocardium by mining publicly available single cell RNA sequencing data from mice receiving HFD/L-NAME compared to STD chow (14). We identified all major cardiac non-myocyte cell types (Figure S3A and S3B) and found fibroblasts as the dominant source of lysyl oxidases in the heart (Figure 3A). Next, we subclustered the isolated CFB population (Figure S3C) and found that Cluster 0, which made up 35% of the CFB population in control mice, expanded to 50% in HFpEF mice (Figure 3B and 3C). To better understand the characteristics of this expanded cluster, we performed gene ontology (GO) analysis of genes which were significantly enriched in this cluster (*i.e.* its marker genes). The term *peptidyl lysine oxidation* stood out by being 50-fold overrepresented within the cluster 0 marker genes (Figure 3D). Consequently, we found higher lysyl oxidase expression across the entire CFB population in HFpEF mice compared to control mice (Figure 3E). Based on the findings that lysyl oxidase family member *LOXL3* correlated significantly with *CD4* (Figure 2I and 2J) and that T cells are causally involved in the development of ECM stiffening (Figure 2E and 2F), we next assessed the abundance of LOXL3 protein in LV tissue of mice after 3-5 weeks of HFD/L-NAME treatment. In line with the timeline of CD4^+^ T cell infiltration and ECM stiffening, the abundance of LOXL3 was significantly higher after 5 weeks, but not 3 weeks of HFD/L-NAME treatment compared to STD controls (Figure 3F). These data support that CFB are a main source of *Loxl3* in the onset of HFD/L-NAME-induced ECM remodeling when CD4^+^ T cells are present in the LV and prompted us to further investigate the relevance of this axis in myocardial stiffening and diastolic dysfunction.

**Figure 3:**
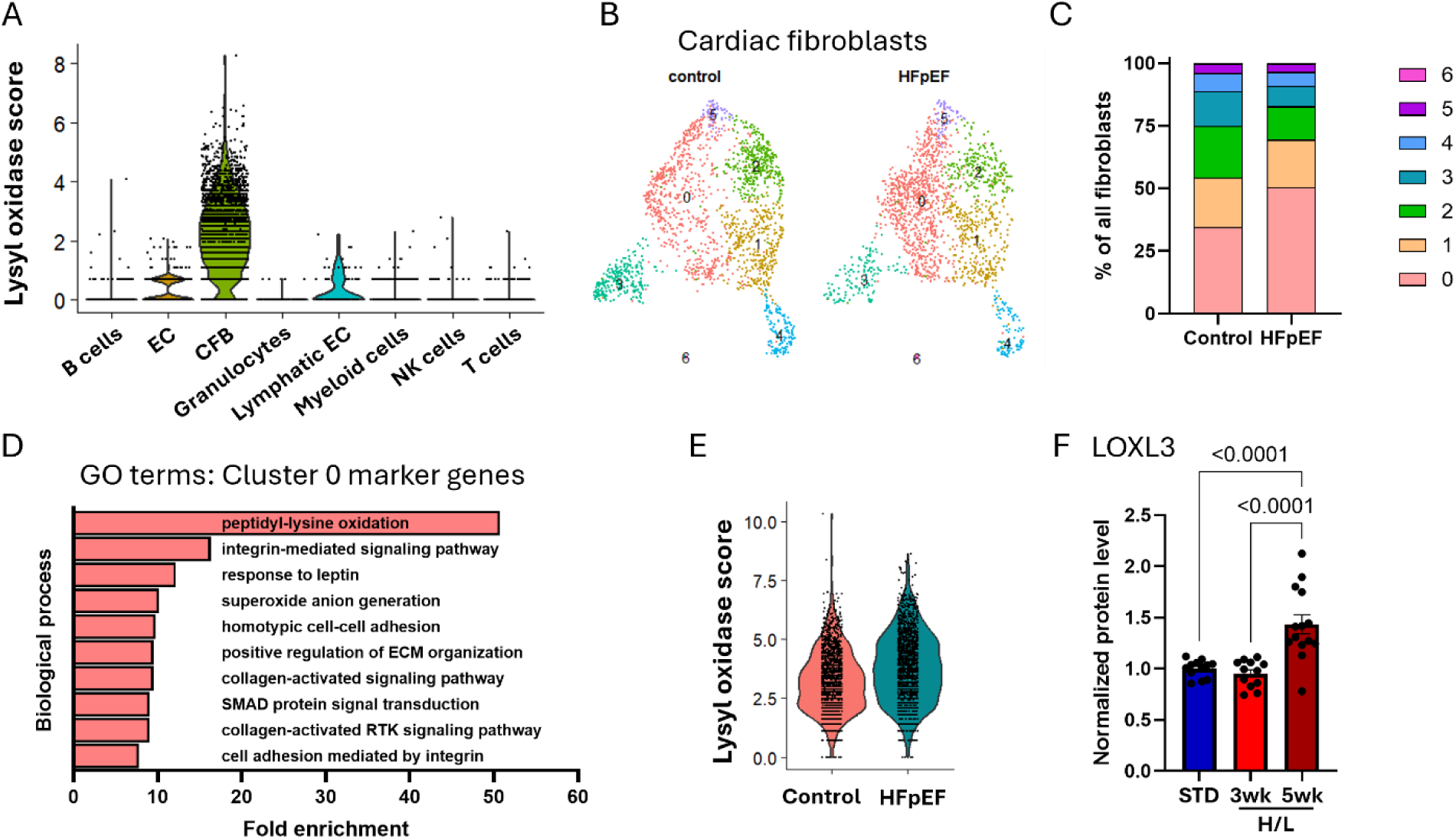
Cardiac fibroblasts express higher levels of lysyl oxidases in cardiometabolic HFpEF. A: Lysyl oxidase expression score across all identified cell types including data from STD-and H/L-treated mice. B: UMAP of CFB isolated from the full data set reveals 6 distinct CFB subclusters. C: Relative abundance of each CFB subcluster in STD-compared to H/L-treated mice. D: Gene ontology (GO) analysis of CFB-cluster 0 marker genes. Shown are the most significantly enriched biological processes. E: Lysyl oxidase expression score across CFB in STD vs H/L-treated mice. F: Cardiac LOXL3 protein levels assessed by ELISA in wild-type C57Bl/6J mice after 3 or 5 weeks of STD- or H/L-treatment (N=12/group).

### CD4^+^ T cell-derived IFNγ induces LOXL3 expression in CFB through HIF1α which causes ECM stiffening in vitro

To gain insight into the mechanism by which CD4^+^ T cells could stimulate *Loxl3* expression in CFB, we next investigated whether the secretome derived from CD4^+^ T cells was able to induce *Loxl3* expression in CFB *in vitro*. We treated primary murine CFBs with the secretome collected from activated CD4^+^ T cell blasts (Figure 4A). After 24h of treatment, the expression of *Loxl3* was significantly higher in CFB treated with the CD4^+^ T cell secretome compared to non-treated control CFB (NC, Figure 4B). Consequently, lysyl oxidase activity in the culture media from CFB was significantly elevated after treatment with CD4^+^ T cell secretome. This demonstrates that, in addition to transcriptional *Loxl3* upregulation, higher levels of active LOXL3 are secreted by CFB in response to CD4^+^ T cell secretome treatment (Figure 4C). We also found that the CD4^+^ T cell secretome did not alter the protein levels of COL1A1 (Figure S4B and S4E), in line with the absence of significant collagen deposition we observed *in vivo*. Thus, we next investigated whether the CD4^+^ T cell secretome increases the ability of CFB to remodel existing collagen instead of producing collagen *de novo*. We first measured the contraction of collagen by CFB disks *in vitro* in the presence or absence of CD4^+^ T cell secretome using TGFβ treatment as a positive control. Treatment of CFB with CD4^+^ secretome resulted in higher collagen gel contraction by CFB compared to non-treated CFB, albeit not to the same extent as observed in TGFβ treated CFB (Figure 4D). We also evaluated the expression levels of the contractile protein αSMA in CFB and found that gene and protein expression were enhanced by TGFβ and not by the activated CD4^+^ T cell secretome, in line with the stronger disc contraction observed by TGFβ compared to the T cell secretome treatment (Figure S4A and S4D). Next, we collected the conditioned media from CFB treated with CD4^+^ T cell secretome (containing active LOXL3) and treated native cardiac dECM to determine its effect on ECM stiffness by uniaxial tensile testing (Figure 4E). Conditioned media from CFB previously exposed to the CD4^+^ T cell secretome resulted in increased stiffness of dECM, compared to dECM treated with control CFB conditioned media (not pre-treated with CD4^+^ T cell secretome and therefore low in LOXL3). This effect was completely abolished in the presence of the lysyl oxidase inhibitor BAPN (Figure 4E and 4F). Taken together, our results demonstrate that a paracrine mediator derived from CD4^+^ T cells induces the expression and secretion of LOXL3 in CFB which subsequently increases the stiffness of native cardiac ECM.

**Figure 4:**
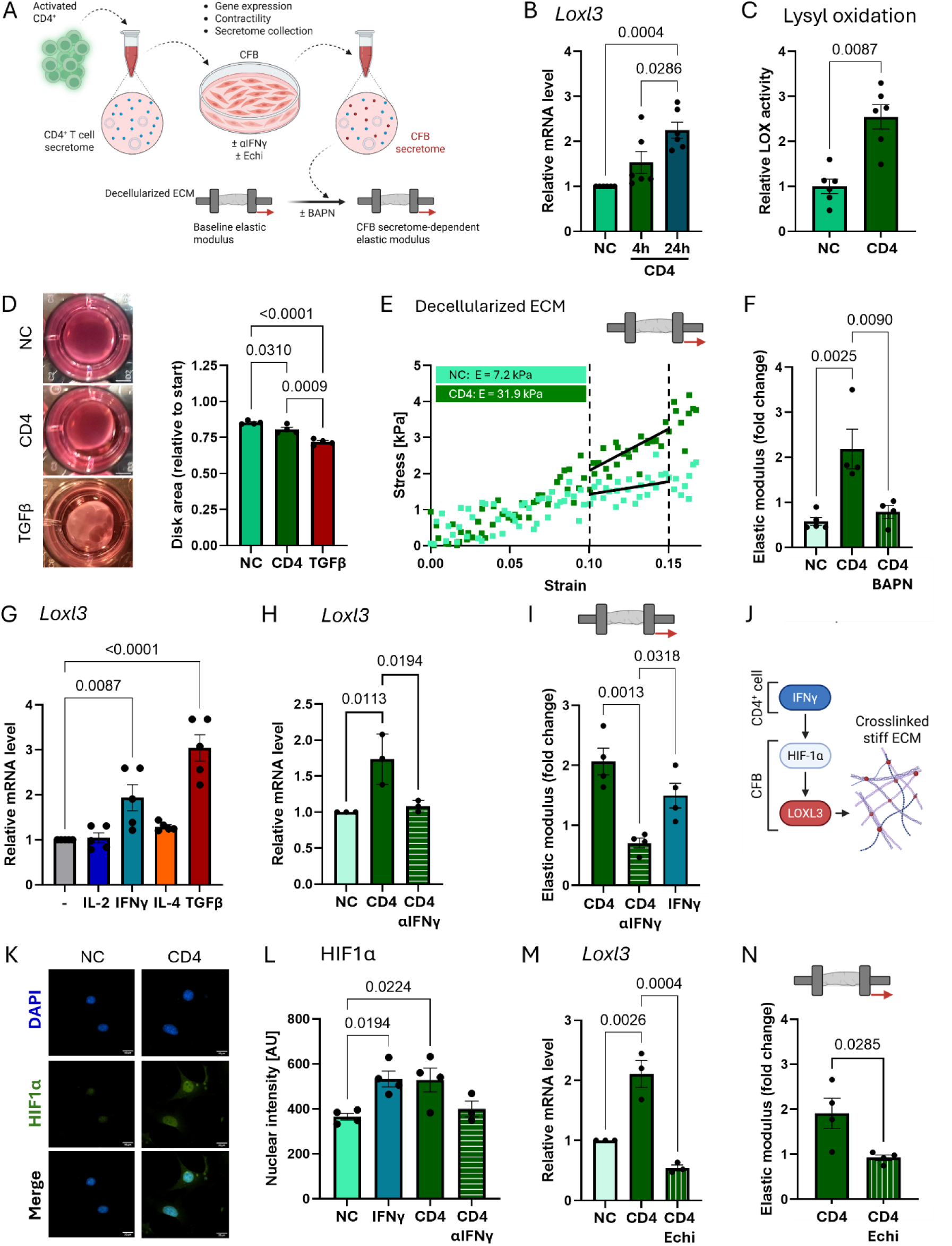
CD4^+^ T cell-derived IFNγ induces LOXL3 expression in CFB through HIF1α which is sufficient for ECM stiffening *in vitro*. A: Cardiac fibroblasts (CFB) isolated from C57Bl/6J mice were treated with the secretome of activated CD4^+^ T cells. B: *Loxl3* mRNA expression levels in CFB treated with the secretome of activated CD4^+^ T cells for the indicated durations (expressed as fold change compared to non-treated CFB (NC), N=6). C: Relative lysyl oxidase activity in culture media from CFB after treatment with CD4^+^ T cell secretome for 24 h (N=6). D: *In vitro* collagen gel contraction by CFB treated with control (NC), CD4^+^ T cell secretome or recombinant TGFβ (100 ng/mL) after 24 h (N=4). E+F: Uniaxial tensile testing of dECM after incubation with CFB secretome. Representative recordings (E) and fold change of Elastic modulus (F) of dECM preparations after incubation with CFB secretome after the indicated treatments in presence or absence of β-aminopropionitrile (BAPN, 500 uM, N=4). G+H: Relative *Loxl3* mRNA levels in CFB after 24 h of treatment with the indicated recombinant cytokines (IL-2: 25 U/mL, IFNγ: 100 U/mL, IL-4: 500 ng/mL, N=5, G) or CD4^+^ T cell secretome in presence or absence of αIFNγ (10 µg/mL, N=3, H) for 24 h. I: Fold change of the Elastic modulus of dECM after incubation with secretome of CFB subjected to the indicated treatments. J: CD4^+^ T cell derived IFNγ triggers HIF1α signaling in CFB to induce the expression LOXL3 and subsequent ECM remodeling. K: Nucleus (DAPI, blue) and HIF1α (green) staining in CFB after 24 h of treatment with CD4^+^ secretome. L: Quantification of the nuclear HIF1α signal intensity from K (N=3-4). M: Relative *Loxl3* mRNA levels after 24 h of treatment with CD4^+^ secretome in the presence or absence of HIF1α inhibitor Echinomycin (Echi, 5 nM, N=3). N: Fold change of the Elastic modulus of dECM after incubation with secretome of CFB treated as indicated (N=4).

To identify the CD4^+^ T cell derived signal that causes the upregulation of LOXL3 in CFB, we treated CFB with cytokines which are well-known to be produced at high levels by CD4^+^ T cells (activated by TCR stimulation with αCD3 and co-stimulation with αCD28 in culture but not polarized towards a specific subset) in culture. TGFβ was used as a profibrotic signal for CFB and positive control. We found that IFNγ induced *Loxl3* expression, whereas IL-2 or IL-4 had no significant effect (Figure 4G). Moreover, neutralizing IFNγ with a selective αIFNγ neutralizing antibody in the CD4^+^ T cell secretome resulted in abrogation of *Loxl3* expression by CFB (αIFNγ, Figure 4H). The ECM stiffening ability of CFB-derived conditioned media was blunted when IFNγ was neutralized during treatment with the CD4^+^ T cell secretome, whereas IFNγ treatment alone reconstituted ECM stiffening (Figure 4I). Taken together, these results demonstrate the requirement of CD4^+^ T cell IFNγ for *Loxl3* expression in CFB and subsequent ECM stiffening by CFB *in vitro*.

To understand the mechanism by which IFNγ induces *Loxl3* expression in CFB, we focused on HIF1α signaling, as lysyl oxidases have been shown to be induced by hypoxia in cancer-associated fibroblasts (18) (Figure 4J). Immunocytochemistry revealed that the CD4^+^ T cell secretome increased the nuclear abundance of HIF1α in CFB. Moreover, HIF1α nuclear localization was dependent on IFNγ, as it was abrogated when IFNγ was neutralized in the secretome (Figure 4K and 4L), demonstrating that CD4^+^ T cell derived IFNγ drives HIF1α nuclear localization in CFB. To further test the requirement of HIF1α in LoxL3 expression and ECM stiffening, we performed similar studies in the presence of the HIF1α inhibitor Echinomycin (Echi). Echi completely abrogated *Loxl3* induction by the CD4^+^ T cell secretome (Figure 4M), as well as ECM stiffening (Figure 4N).

Taken together, these data demonstrate that CD4^+^ T cell IFNγ is necessary for HIF1α nuclear localization and subsequent *Loxl3* expression in CFB, and that inhibition of HIF1α or IFNγ prevent ECM stiffening induced by T cells.

### IFNγ and lysyl oxidation are required for mechanical ECM remodeling and diastolic dysfunction in cardiometabolic HFpEF

To assess the relevance of IFNγ-mediated HIF1α activation and subsequent LOXL3 upregulation *in vivo*, WT mice were injected with recombinant IFNγ for 5 consecutive days (Figure 5A). We found higher levels of *Hif1a* and *Loxl3* mRNA (Figure 5B and 5C) as well as LOXL3 protein (Figure 5D) in whole LV lysates of IFNγ-treated mice compared to PBS-treated controls. Consequently, we asked if the absence of IFNγ would change the pathological outcome of HFD/L-NAME treatment *in vivo*. We treated IFNγ-deficient (*Ifng^−/−^*) mice alongside WT mice with HFD/L-NAME for 5 weeks (Figure 5E). Strikingly, the LV-ECM of *Ifng^−/−^* mice was significantly softer than that of WT mice, as determined by lower elastic modulus (Figure 5F). Moreover, *Ifng^−/−^*mice were protected from diastolic dysfunction in response to HFD/L-NAME, supporting the requirement of IFNγ for ECM stiffening and diastolic dysfunction in cardiometabolic HFpEF (Figure 5G).

**Figure 5:**
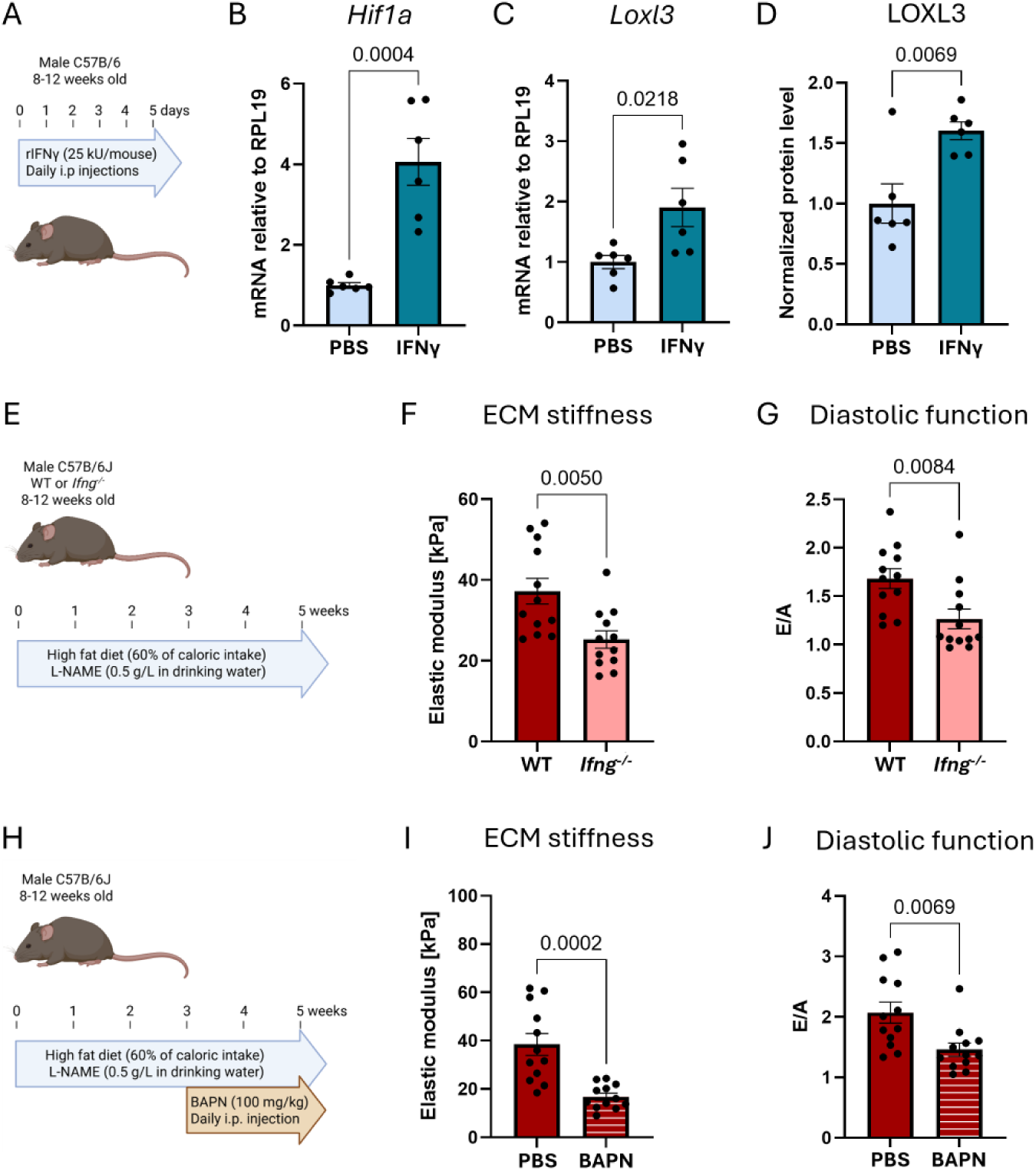
*In vivo*, IFNγ and lysyl oxidation are required for ECM stiffening and diastolic dysfunction. A: Wild-type C57Bl/6J mice received 25 kU recombinant IFNγ or PBS for 5 consecutive days by i.p. injection (N=6). B-C: Relative *Hif1α* (B) and *Loxl3* (C) mRNA as well as LOXL3 protein (D) levels in the LV of PBS or IFNγ treated mice. E: Wild-type or *Ifng^−/−^* C57Bl/6J mice were subjected to H/L treatment for 5 weeks. F-G: Elastic modulus of LV-ECM (F) and diastolic function expressed as E/A ratio (G) in wild-type and *Ifng^−/−^* mice after 5 weeks of HFD/L-NAME treatment (N=12/group). H: Wild-type C57Bl/6J were subjected to HFD/L-NAME treatment for 5 weeks with daily PBS or β-aminopropionitrile (BAPN, 100 mg/kg) injections from week 3 to 5. I-J: Elastic modulus of LV-ECM (I) and diastolic function expressed as E/A ratio (H) in wild-type mice receiving PBS or BAPN during weeks 3-5 of HFD/L-NAME treatment (N=12/group).

To investigate the importance of lysyl oxidase-mediated ECM stiffening for diastolic dysfunction in response to HFD/L-NAME *in vivo*, we treated WT mice with BAPN, starting 3 weeks after switching their diet to HFD/L-NAME (*i.e.* prior to CD4^+^ T cell infiltration (Figure 2A), ECM stiffening (Figure 2B) and cardiac LOXL3 upregulation (Figure 3F)) and continued until the end of the experiment at 5 weeks of HFD/L-NAME (Figure 5H). Strikingly, BAPN treatment resulted in significantly lower ECM stiffness in BAPN treated mice compared to PBS treated mice (Figure 5I). Consequently, BAPN treated mice did not develop significant diastolic dysfunction (Figure 5J). Taken together, we show that IFNγ is sufficient to induce cardiac LOXL3 expression *in vivo*. Both, IFNγ and lysyl oxidation are required for cardiac stiffening and diastolic dysfunction in response to HFD/L-NAME.

## Discussion

Here, we investigated the role of mechanical ECM remodeling in the pathology of cardiometabolic HFpEF and assessed the contribution of CD4⁺ T cells in steering mechanical ECM remodeling through collagen crosslinking. We identified elevated ECM stiffness as a crucial contributor to diastolic dysfunction and describe a novel mechanism of T cell IFNγ instructed upregulation of LOXL3 in CFB that drives this stiffening. Cardiac tissue fibrosis, defined as excessive accumulation of ECM, is associated with the majority of cardiac diseases. In HFpEF patients the occurrence of fibrosis appears to be variable. While a number of studies show mild but significant fibrosis in patients as well as animal models (17,19,20), other studies suggest that fibrosis plays a minor role in regard to diastolic function. In fact, HFpEF patients with the best-preserved ejection fractions may be the least likely to show significant fibrosis (21). Our results demonstrate that enhanced crosslinking of ECM by lysyl oxidases results in ECM stiffening in experimental cardiometabolic HFpEF in the absence of histologically detectable fibrosis. Pharmacological inhibition of lysyl oxidases prevented both ECM stiffening and diastolic dysfunction. This supports that enhanced ECM crosslinking is sufficient to cause diastolic dysfunction and does not require *de novo* ECM accumulation. This is highlighted by our functional *in vitro* studies, in which LOXL3 enrichment in culture media was solely responsible for the stiffening of native cardiac ECM. Using mass spectrometry based in-depth analysis of the cardiac ECM, we previously identified sub-histological increases of ECM deposition in the LV of mice with hypertensive HFpEF, a phenomenon termed hidden fibrosis (22). Our data in cardiometabolic HFpEF supports the conclusion that ECM remodeling does not need to be visible as collagen deposition to be functionally relevant. In fact, our data collected after only 5 weeks of HFD/L-NAME treatment suggests that sub-histological mechanical remodeling may precede the development of prominent fibrosis characterized by collagen deposition.

We demonstrate the causal involvement of CD4^+^ T cells in mechanical ECM remodeling in HFD/L-NAME-induced cardiometabolic HFpEF. This is in line with our previous work which identified systemic CD4^+^ T cell activation and CD4^+^ T cell cardiotropism as crucial features of cardiometabolic HFpEF (5). Sequencing data from human HFpEF patients suggests a strong correlation between the presence of CD4^+^ cells and the expression of *LOXL3*. While we did not observe a significant difference in *LOXL3* expression between control and HFpEF patients, the correlation between *CD4* and *LOXL3* was statistically significant across groups suggesting that the abundance of CD4^+^ T cells in patient tissue relates to the extent of collagen crosslinking through LOXL3. The lysyl oxidase LOXL2 has emerged as a promising therapeutic target in HF patients with hypertension and aortic stenosis (24). Our work suggests that targeting LOXL3 may serve as a candidate novel therapeutic strategy in cardiometabolic HFpEF patients.

Mechanistically, we discovered a novel, contact-independent communication pathway between cardiac infiltrated CD4^+^ T cells and CFB, mediated by the canonical Th1 cytokine IFNγ. Our in vitro studies demonstrate that T cell derived IFNγ is required for CFB expression of LOXL3, and our *in vivo* studies using recombinant IFNγ and *Ifng^−/−^* mice support this and its importance for diastolic function. However, *in vivo*, T cells are not the exclusive source of IFNγ in the heart as it is also produced by natural killer (NK) and NKT cells. NKT cells may be of particular interest in the context of hyperlipidemia as they are activated by lipid antigens presented by the non-classical MHC-I-like molecule CD1d (23). Whether IFNγ-producing NK(T) cells are also enriched in the myocardium and how they may contribute to T cell-CFB communication and ECM crosslinking through LOXL3 in this model will be the subject of future studies.

The effects IFNγ on CFB are manifold. Our data paint a complex picture, in which on the one hand, IFNγ upregulates *Loxl3* similarly to TGFβ, and on the other hand does not enhance the fibrotic marker genes *Acta2* and *Col1a1*. Even though seemingly opposing, both effects are supported by previous studies. Our lab demonstrated that the integrin α4-dependent interaction between IFNγ^+^ Th1 cells and CFB leads to TGFβ production and subsequent transformation of CFB into fibrotic myofibroblasts in pressure overload-induced HF (7). Others showed that IFNγ counteracts the effects of TGFß in CFB and favors an inflammatory phenotype over a fibrotic phenotype (23). Altogether, this highlights the complexity of the CFB phenotype and the need for delineating CFB responses to stimulation in a disease-specific context. CFB heterogeneity and disease-specificity have been explored using single cell sequencing and computational approaches to compare their transcriptomic signatures in multiple murine HF models. In line with our data showing the requirement for HIF1α for IFNγ-driven LOXL3 expression, single cell sequencing identified a hypoxia signaling signature as specific feature of HFpEF fibroblasts compared to those from HFrEF models like transverse aortic constriction or long term remodeling after myocardial infarction (14).

While our work demonstrates that mechanical ECM remodeling in cardiometabolic HFpEF is triggered by CD4^+^ T cell derived IFNγ and executed by CFB releasing LOXL3, there are some limitations that need to be acknowledged. Our *in vivo* studies have been performed in male mice as female mice are more resistant to diastolic dysfunction in the 2-hit model of cardiometabolic HFpEF even up to 15 weeks of HFD/L-NAME (25). However, the IFNγ-LOXL3 axis we report herein for CD4^+^ T cell-CFB crosstalk may show sex specific differences. Further, we focused our current study on the HFD/L-NAME HFpEF model, so the relevance of our findings to other models of HFpEF, and therefore potentially to other patient subgroups, remains untested. In a hypertensive HFpEF model using DOCA-salt treatment and uninephrectomy, we also found that ECM enrichment was not detectable by histology (22). Whether such small increases of ECM deposition alone are sufficient to cause diastolic dysfunction or whether additional crosslinking such as through the mechanisms described here are at play requires further studies. The IFNγ-mediated transcriptional upregulation of LOXL3 we describe here may involve changes in chromatin accessibility, consistent with the mechanism we described in the hypertensive HFpEF model (22). In addition to T cells, other immune cell types including macrophages are important contributors to pathology in the 2-hit model of cardiometabolic HFpEF (26), and it is possible that T cell-macrophage interactions also take place. Lastly, beyond the mechanical properties of the ECM, cardiomyocyte relaxation is a critical contributor to diastolic function. Our previous study suggests that impaired cardiomyocyte relaxation is T cell dependent (5). However, how T cells, CFB and potentially macrophages interact with cardiomyocytes in the context of cardiometabolic HFpEF and contribute to ECM mechanical remodeling remains to be explored.

Taken together, our work identifies a previously unrecognized immune-stromal axis by which T cell inflammation directly affects mechanical ECM remodeling and diastolic dysfunction in cardiometabolic HFpEF. We demonstrate a novel mechanism of IFNγ-dependent communication between cardiac infiltrated CD4^+^ T cells and CFB and establish a novel mechanistic link between adaptive immunity and cardiac mechanics in HFpEF. The resulting increase in LOXL3 expression and collagen crosslinking is a crucial driver of diastolic dysfunction in cardiometabolic HFpEF. This novel immune-stromal interaction reveals potential new targets to mitigate cardiac stiffening and diastolic dysfunction in cardiometabolic HFpEF.

## Novelty and Significance

### What is known?

- Stiffness of the extracellular matrix is an important determinant of diastolic function
- Cardiac fibroblasts are the major cell type responsible for cardiac extracellular matrix homeostasis and remodeling, and are sensitive to cytokine stimulation
- CD4^+^ T cells infiltrate the myocardium and are required for diastolic dysfunction in cardiometabolic HFpEF

### What new information does this article contribute?

- Enhanced crosslinking by lysyl oxidation causes diastolic dysfunction in the absence of histological fibrosis in cardiometabolic HFpEF
- CD4^+^ T cell-derived IFNγ drives the expression of the lysyl oxidase LOXL3 in cardiac fibroblasts through HIF1α
- Interference with this axis by pharmacologic inhibition of lysyl oxidation or genetic knock-out of IFNγ protects mice from pathologic ECM stiffening and diastolic dysfunction

Rising prevalence and limited treatment options render heart failure with preserved ejection fraction (HFpEF) one of the biggest unmet needs of modern medicine. Our work demonstrates a novel mechanism by which cardiac CD4^+^ T cells interact with cardiac fibroblasts to increase extracellular matrix stiffness and cause diastolic dysfunction in experimental cardiometabolic HFpEF. This suggests the lysyl oxidase LOXL3 and the inflammatory cytokine IFNγ as novel therapeutic targets for patients with cardiometabolic HFpEF.

## Supplemental Material

Tables S1-6

Figures S1-4

## Non-standard Abbreviations and Acronyms

BAPN: ß-aminopropionitrile
CFB: Cardiac fibroblast
Echi: Echinomycin
ECM: Extracellular matrix
E/A: Early/atrial mitral valve inflow velocity
HFD: High fat diet
HFpEF: Heart failure with preserved ejection fraction
HIF1α: Hypoxia-inducible factor-1α
IFNγ: Interferon-γ
LOXL3: Lysyl oxidase-like 3
LV: Left ventricle
L-NAME: L-nitro-arginine-methyl ester
STD: Standard diet
TGFß: Transforming growth factor ß

