## Supplemental information for "T cell-derived IFNγ instructs ECM crosslinking by cardiac fibroblasts through LOXL3 in experimental cardiometabolic HFpEF"

**Running title: T cell-instructed ECM remodeling in HFpEF**

*Ramona Emig PhD[1,2], Zachary Robbe[2], Celina Kley[3], Sasha Smolgovsky PhD[2,4], Joshua G Travers PhD[5], Robert M Blanton MD[6], Timothy A McKinsey PhD[5], Lauren D Black PhD[7], Pilar Alcaide PhD[1,2]*

[1]Department of Microbiology and Immunology, Miller School of Medicine, University of Miami, Miami, FL, USA
[2]Department of Immunology, School of Medicine, Tufts University, Boston, MA, USA
[3]German Centre of Cardiovascular Research, University of Heidelberg, Heidelberg, Germany
[4]Department of Laboratory Medicine and Pathology, University of Washington, Seattle, WA, USA
[5]Department of Medicine, Division of Cardiology, and Consortium for Fibrosis Research & Translation, University of Colorado Anschutz Medical Campus, Aurora, CO, USA
[6]Molecular Cardiology Research Institute, School of Medicine, Tufts University, Boston, USA
[7]Department for Bioengineering, Tufts University, Boston, MA, USA

### Supplemental figures

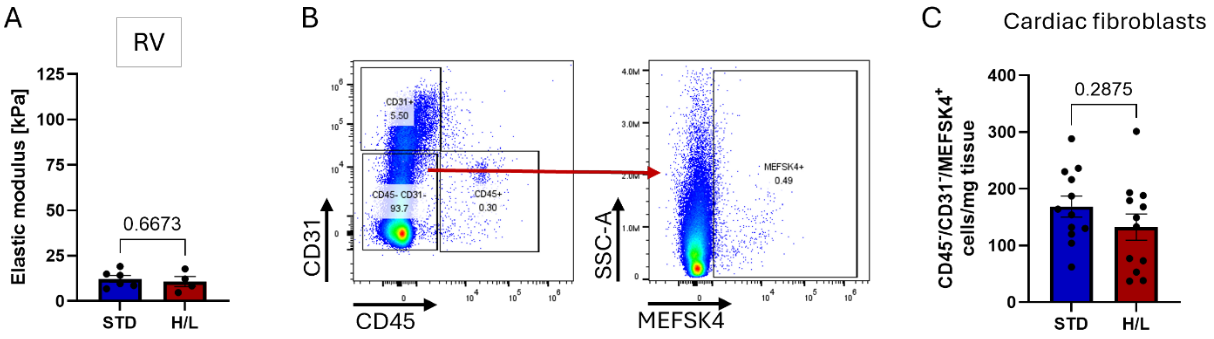


**Figure S1:** A: Elastic modulus of right ventricular ECM samples from C57Bl/6J mice subjected to normal diet (STD) or high fat diet and L-NAME (H/L, N=12/group). B: Gating strategy for flow cytometry-based quantification of left ventricular $CD45^-/CD31^-/MEFSK4^+$ fibroblasts. C: Numbers of left ventricular fibroblasts per mg tissue from STD or H/L treated mice (N=12/group).

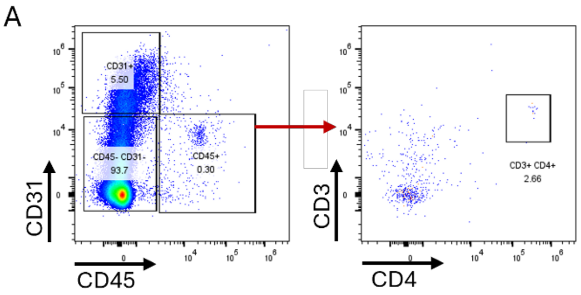


**Figure S2:** A: Gating strategy for flow cytometry-based quantification of left ventricular $CD3^+/CD4^+$ T cells.

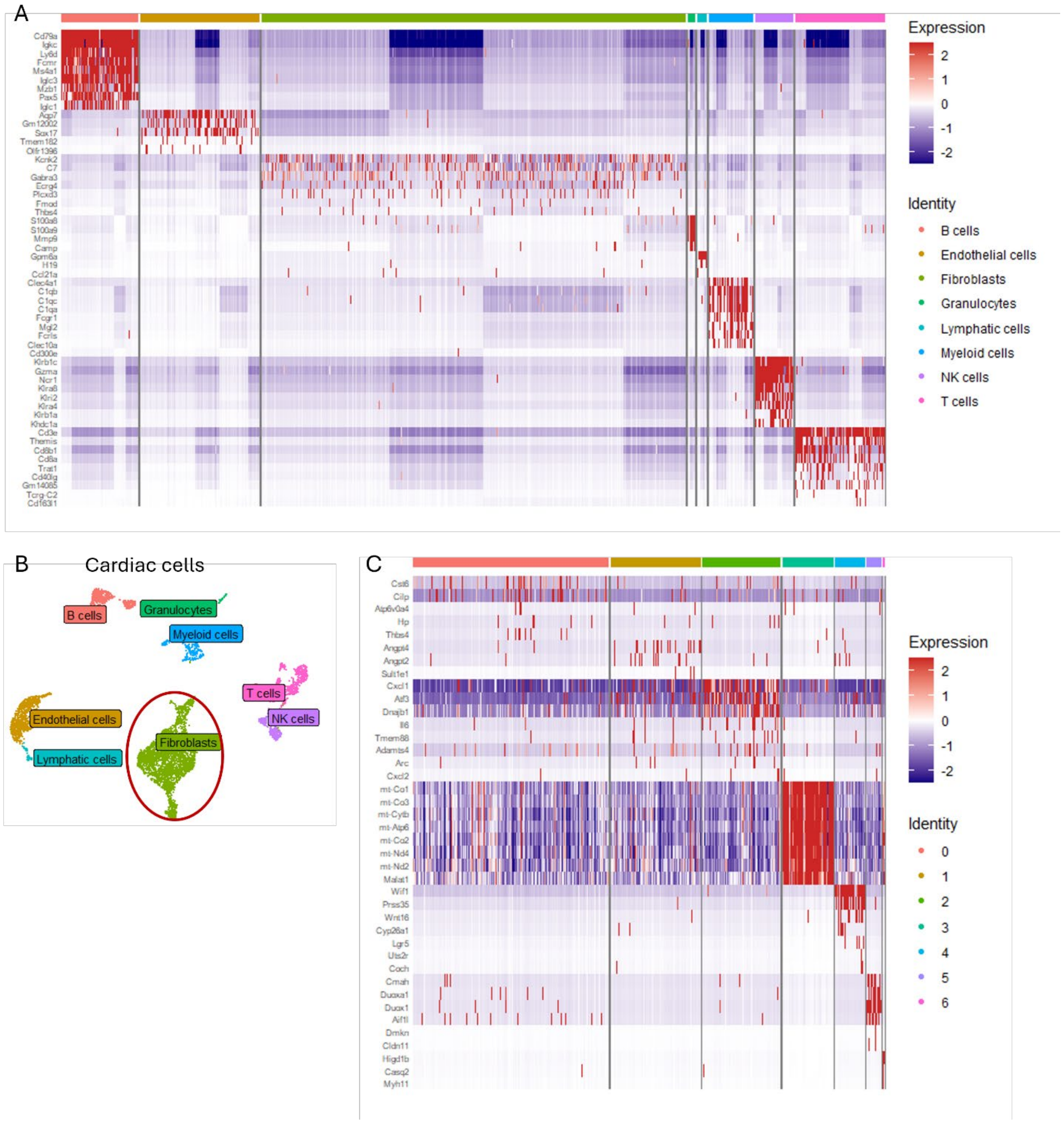


**Figure S3:** Single cell RNA sequencing analysis of left ventricular non-myocytes from mice subjected to standard diet or high fat diet+L-NAME. A: Heatmap of the top marker genes for each identified cell type. B: UMAP showing that single cell RNA sequencing of murine cardiac non-myocytes reveals all major cardiac non-myocyte cell populations and allows identification of cardiac fibroblasts. C: Heatmap plot of the Top 10 marker genes for each identified cluster of cardiac fibroblasts.

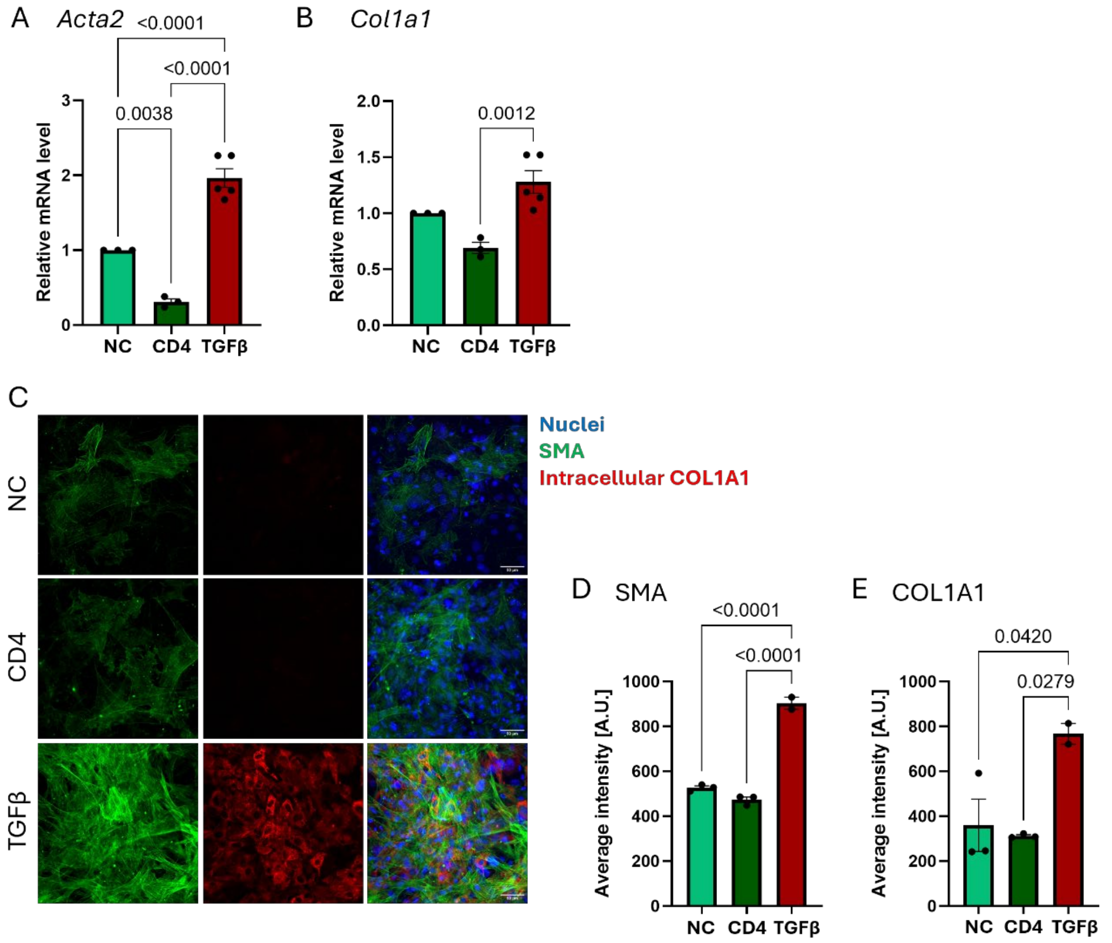


**Figure S4:** A-B: Relative mRNA *Acta2* (A) and *Col1a1* (B) level in cardiac fibroblasts after 24h of treatment with CD4$^+$ T cell secretome or recombinant TGFß (100 ng/mL, N=3-5). C: Immunocytochemistry for cardiac fibroblast nuclei (DAPI, blue), smooth muscle actin (SMA, green) or Collagen-1a1 (COL1A1, red) after 24 h of treatment with CD4$^+$ T cell secretome or recombinant TGFß (100 ng/mL). D-E: Quantification of fluorescence intensity of SMA (D) or COL1A1 (E) staining. Each data point represents the average of 5 images acquired from N=2-3 independent experiments.

### Supplemental tables

**Table S1:** Antibodies used in flow cytometry experiments.

| Target | Fluorophore | Dilution | Company | Catalog no |
|---|---|---|---|---|
| **CD45.2** | BUV-805 | 1:100 | BD Bioscience | 569200 |
| **CD31** | BV-421 | 1:100 | BioLegend | 102424 |
| **CD3** | BV-605 | 1:100 | BioLegend | 100351 |
| **CD4** | PE | 1:100 | BioLegend | 100512 |
| **CD8** | BV-711 | 1:100 | BioLegend | 100748 |
| **MEFSK4** | APC | 1:50 | Miltenyi Biotech | 105714 |
| **CD16/CD32 (Fc block)** | - | 1:50 | BioLegend | 101320 |

**Table S2:** Primer sequences (5’ to 3’) used in quantitative real-time PCR experiments.

| Target gene | Forward primer | Reverse primer |
|---|---|---|
| ***Loxl3*** | CTACTGCTGCTACACTGTCTGT | GTAGTGAATGTTGCATCTCACCA |
| ***Acta2*** | GTCCCAGACATCAGGGAGTAA | TCGGATACTTCAGCGTCAGGA |
| ***Col1a1*** | GCTCCTCTTAGGGGCCACT | ATTGGGGACCCTTAGGCCAT |
| ***Hif1a*** | ACCTTCATCGGAAACTCCAAAG | CTGTTAGGCTGGGAAAAGTTAGG |

**Table S3:** Male C57Bl/6J mice were fed a standard diet or combination of high fat diet (60% fat) and L-NAME (0.5 g/L in drinking water) for 5 weeks. Cardiac function and dimensions were assessed by echocardiography. Data is presented as mean±standard error to the mean (N=12).

| | **STD** | **H/L** | **p-value** |
|---|---|---|---|
| **Body weight (g)** | 26.18±0.29 | 32.87±0.72 | **<0.0001** |
| **Ejection fraction (%)** | 57.04±1.72 | 58.47±0.75 | 0.4726 |
| **Anterior wall thickness systole (mm)** | 1.24±0.02 | 1.39±0.03 | **0.0012** |
| **Anterior wall thickness diastole (mm)** | 0.86±0.02 | 0.97±0.03 | **0.0060** |
| **Internal diastolic diameter (mm)** | 4.16±0.06 | 3.67±0.09 | **0.0002** |
| **Internal systolic diameter (mm)** | 2.96±0.08 | 2.59±0.04 | **0.0004** |
| **E/A** | 1.23±0.08 | 1.56±0.06 | **<0.0001** |
| **E/E’** | -21.69±0.40 | -24.08±1.05 | 0.0531 |

**Table S4:** Male wildtype and *Tcra*$^{-/-}$ C57Bl/6J mice were fed a standard diet or combination of high fat diet (60% fat) and L-NAME (0.5 g/L in drinking water) for 5 weeks. Cardiac function and dimensions were assessed by echocardiography. Data is presented as mean±standard error to the mean (N=6-7). ****: p<0.0001 compared to STD, $^{\#\#\#\#}$: p<0.0001 compared to wildtype.

| | **Wildtype STD** | **Wildtype H/L** | ***Tcra*$^{-/-}$ STD** | ***Tcra*$^{-/-}$ H/L** |
|---|---|---|---|---|
| **N** | 6 | 6 | 7 | 8 |
| **Body weight (g)** | 26.07±0.48 | 32.88±1.05**** | 25.87±0.55 | 34.09±0.95**** |
| **Ejection fraction (%)** | 55.79±2.76 | 59.08±1.04 | 58.00±5.33 | 54.13±5.46 |
| **Anterior wall thickness systole (mm)** | 1.24±0.03 | 1.38±0.05 | 0.96±0.06 | 1.22±0.06 |
| **Anterior wall thickness diastole (mm)** | 0.86±0.02 | 0.98±0.05 | 0.84±0.07 | 0.99±0.11 |
| **Internal diastolic diameter (mm)** | 4.17±0.09 | 3.72±0.06 | 3.82±0.08 | 3.81±0.19 |
| **Internal systolic diameter (mm)** | 2.96±0.11 | 2.58±0.06 | 2.65±0.09 | 2.73±0.14 |
| **E/A** | 1.22±0.04 | 1.57±0.06** | 1.26±0.07 | 1.3±0.06$^{\#\#}$ |
| **E/E’** | -21.86±0.58 | -24.17±1.50 | -31.43±3.01 | -35.75±9.34 |

**Table S5:** Male wildtype and *Ifng*$^{-/-}$ C57Bl/6J mice were fed the combination of high fat diet (60% fat) and L-NAME (0.5 g/L in drinking water) for 5 weeks. Cardiac function and dimensions were assessed by echocardiography. Data is presented as mean±standard error to the mean (N=12).

| | **Wildtype** | ***Ifng*$^{-/-}$** | **p-value** |
|---|---|---|---|
| **Body weight (g)** | 33.35±1.24 | 33.19±0.64 | 0.9146 |
| **LV weight/Tibia length (mg/mm)** | 5.82±0.09 | 5.34±0.07 | **0.0008** |
| **Ejection fraction (%)** | 65.02±2.53 | 67.20±2.30 | 0.5510 |
| **Anterior wall thickness systole (mm)** | 1.55±0.06 | 1.54±0.04 | 0.9630 |
| **Anterior wall thickness diastole (mm)** | 1.13±0.05 | 1.08±0.05 | 0.5207 |
| **Internal diastolic diameter (mm)** | 3.58±0.07 | 3.48±0.07 | 0.3500 |
| **Internal systolic diameter (mm)** | 2.32±0.10 | 2.20±0.08 | 0.3733 |
| **E/A** | 1.68±0.10 | 1.27±010 | **0.0084** |
| **E/E'** | -36.01±4.54 | -34.83±2.52 | 0.8334 |

**Table S6:** Male C57Bl/6J mice were fed the combination of high fat diet (60% fat) and L-NAME (0.5 g/L in drinking water) for 5 weeks. During weeks 3-5, mice received daily i.p. injections of PBS or ß-aminopropionitrile (BAPN, 100 mg/mL in PBS). Cardiac function and dimensions were assessed by echocardiography. Data is presented as mean±standard error to the mean (N=12).

| | **PBS** | **BAPN** | **p-value** |
|---|---|---|---|
| **Body weight (g)** | 32.43±0.89 | 30.81±0.72 | 0.1880 |
| **LV weight/Tibia length (mg/mm)** | 4.83±0.13 | 4.73±0.09 | 0.5751 |
| **Ejection fraction (%)** | 55.95±2.29 | 59.75±2.28 | 0.2750 |
| **Anterior wall thickness systole (mm)** | 1.37±0.04 | 1.44±0.04 | 0.3095 |
| **Anterior wall thickness diastole (mm)** | 0.95±0.03 | 1.02±0.03 | 0.1684 |
| **Internal diastolic diameter (mm)** | 3.75±0.10 | 3.68±0.04 | 0.4994 |
| **Internal systolic diameter (mm)** | 2.67±0.11 | 2.51±0.07 | 0.2189 |
| **E/A** | 2.07±0.17 | 1.46±0.11 | **0.0069** |
| **E/E'** | -36.61±2.67 | -31.14±1.55 | 0.0988 |
